# In response to Li *et al.*: Linker histones function in *Drosophila* embryogenesis

**DOI:** 10.1101/2020.03.21.001529

**Authors:** Albert Carbonell, Lazslo Henn, Juan Pérez-Roldán, Srividya Tamirisa, Anikó Szabó, Imre M. Boros, Fernando Azorín

## Abstract

In an earlier paper (Pérez-Montero et al., 2013), we reported that the embryonic linker histone of *Drosophila* dBigH1 was essential for early *Drosophila* embryogenesis since embryos homozygous for the *bigH1^100^* mutation showed strong defects and did not survive beyond zygotic genome activation (ZGA) at cellularization. Recent results challenge these observations since null *bigH1* mutations generated by CRISPR/Cas9 methodology turn out to be homozygous viable, as reported in Li *et al*. (2019) and here. In this regard, Li *et al*. described a novel mechanism by which lack of dBigH1 is compensated by the early expression of maternal dH1. Here, we confirm this observation and show that such compensatory mechanism is not activated in *bigH1^100^* embryos.

## INTRODUCTION

Metazoans usually encode multiple histone H1 variants, some of which are specifically expressed in the germline (reviewed in Pérez-Montero et al., 2016). In vertebrates, female and male germline specific H1s are usually different. Female-specific variants accumulate in the oocyte and are retained during early embryogenesis (Pérez-Montero et al., 2016). In *Drosophila*, H1 complexity is low since it encodes for a single ubiquitously expressed somatic dH1 variant and a single germline specific dBigH1 variant (reviewed in Bayona-Feliu et al., 2016). dBigH1 is expressed in both the female and male germline, and it is present in the early embryo until zygotic genome activation (ZGA) at cellularization (Carbonell et al., 2017; Pérez-Montero et al., 2013). At this stage, dBigH1 is replaced by somatic dH1 in somatic cells, whereas it is retained in the primordial germ cells (PGC) (Pérez-Montero et al., 2013). In previous studies, we generated a loss-of-function *bigH1^100^* allele that showed strong embryonic lethality with no homozygous *bigH1^100^* embryos surviving cellularization (Pérez-Montero et al., 2013), suggesting that dBigH1 is essential for early embryo development. Recent results by Li *et al*. challenged this view since null *bigH1* CRISPR alleles were found to be homozygous viable (Li et al., 2019). Notably, Li *et al*. showed that the lack of dBigH1 was compensated by the early expression of maternal dH1 (Li et al., 2019).

## MATERIALS AND METHODS

### Drosophila stocks

*bigH1^100^* allele is described in (Pérez-Montero et al., 2016). *bigH1^NULL^* and *bigH1^NSTOP^* alleles were generated by CRISPR/Cas9 mediated homologous recombination. CRISPR Optimal Target Finder (http://targetfinder.flycrispr.neuro.brown.edu/) was used for identification of gRNA target sites downstream and upstream of the *dBigH1* encoding genomic locus. Upstream (5’-ATTAGCAGTGTTATTCCATA-3’) and downstream (5’-ATAATACCTCTAGAAGGAAT-3’) gRNA sequences were cloned into pCFD4 plasmid (Port et al., 2014) and injected into *y[1] v[1] P{nos-phiC31\int.NLS}X; P{CaryP}attP40* embryos (Bischof et al., 2007) (500ng/μl plasmid DNA in injection buffer (5mM KCl, 0.1mM KH_2_PO_4_, pH:7.8)) to insert transgenic gRNA genes into the 2nd chromosome of the *Drosophila* genome.

A 4640 bp long genomic fragment containing the *dBigH1* locus was amplified with *dBigH1 Rev* (5’-GGACACACTGACATTTAGCTGTTTGG-3’) and *dBigH1 Fw* (5’-TACTCCGTAATTGATGAGATTCCGCC-3’) primers and inserted into pTZ57R/T plasmid (Thermofischer Scientific). On this donor plasmid PAM sequences of upstream and downstream gRNA target sites were mutated by PCR mutagenesis using BigH1 PAM upstream (5’-ACCTTCCGTACGCTCATTTTTAGAATTAACTCCTCTTCTTCTTTAGCAGTGTTAACATATG-3’) and BigH1 PAM downstream (5’-AACTCAATTGAAATGAGAAAATGTGTTTATAATAGCTCTAGAAGGAATAAGCGATAACCGAG-3’) primers. *bigH1^NSTOP^* sequence was generated by PCR mutagenesis using NSTOP primer (5’-CGATTCTGACAACCCCAAGTCGATGGTCTGAAAACCAAAGGG-3’) exchanging P58 with STOP codon and 3xFLAG epitope encoding sequence was built following the first methionine. 3xP3 promoter driven *dsRed* sequence as a marker gene was introduced into *bigH1^NSTOP^* allele carrying donor plasmid, followed by 16bp of the *dBigH1* 5’UTR. *bigH1^NULL^* sequence was generated by Sequence and Ligation Independent Cloning (SLIC) (Jeong et al., 2012). In this case the entire dBigH1 coding sequence was replaced with mCherry sequence surrounded by loxP sites. For SLIC reactions the following primers were used: loxP 5UTR (5’- CTTCGTATAATGTATGCTATACGAAGTTATCATGTTATTAGTTGGAAATTAAATTGAACAAAAATTAGAAATACAACTC-3’), loxP mCh Fw (5’-TCGTATAGCAT ACATTATACGAAGTTATTAGTGAGCAAGGGCGAGGAGGA-3’), loxP 3UTR (5’-TTCGTATAGCATACATTATACGAAGTTATAATTATGATTTATATGTTTTTTTTTCAGTACATGTG-3’), and loxP mCh Rev (5’-ACTTCGTATAATGTATGCTATACGAAGTTATCTTGTACAGCTCGTCCATGCCG-3’). *bigH1^NSTOP^* and *bigH1^NULL^* encoding donor plasmids were injected (500ng/μl plasmid DNA in injection buffer) into embryos laid by *y[1] M{vas-Cas9}ZH-2A w^1118^* females (BDSC_51323) crossed with transgenic *dBigH1* gRNA expressing males. Gene replacements were identified by *dsRed* expression in male descendants of injected males (in the case of *bigH1^NSTOP^* allele) or PCR on genomic DNA using primers specific for mCherry (mCherry Fw: 5’-GTGAAGCTGCGCGGCACC-3’) and *dBigH1* (dBigH1 rev0: 5’-TGTGGAGAATACCTATGACGATTGCG-3’) (in the case of *bigH1^NULL^*) and Sanger sequencing of genomic DNA of both mutants, as well.

### Determination of embryo hatching rate

Homozygous mutant and wild type embryos were collected at 25°C and hatching rates were determined 36h after egg laying by counting hatched and unhatched embryos in three independent replicates of each genotype (50-200 embryos/replicate, 120 on average).

### Antibodies

Rabbit αdH1 (1:8000 for IF) was kindly provided by Dr. J. Kadonaga and is described in (Bayona-Feliu et al., 2017). Rabbit αdBigH1 (1:400 for IF; 1:5000 for WB) is described in (Pérez-Montero et al., 2013). αH3 (1:2000 for WB) was commercially available (Cell Signaling 9715S).

### Immunofluorescence analysis

Immunostaining of embryos was performed according to standard methods. In brief, 0-2h embryos were collected in peach juice plates, dechorionated with 50% bleach for 2 min, fixed for 20 min in 47% heptane and 5.8% formaldehyde, and devitellinized in MeOH. The embryos were then permeabilized in PBS, 0.3% Triton; blocked in PBS, 0.3% Triton, 2% BSA; and incubated at 4°C overnight with the indicated antibodies. For visualization, slides were mounted in Mowiol (Calbiochem-Novabiochem) containing 0.2 ng/μl DAPI (Sigma) and visualized in a confocal microscope (Leica SPE).

### Western blot (WB) analysis

Total protein extracts were obtained by homogenizing 40 ovaries in SDS-PAGE loading buffer. WB was carried out according to standard methods.

## RESULTS

We generated two null *bigH1* CRISPR alleles: *bigH1^NULL^*, in which the complete coding sequence of *bigH1* was replaced by *mCherry*, and *bigH1^NSTOP^*, in which an early stop was generated into the N-terminal domain of dBigH1 (**Figure 1A**) (see **Materials and Methods**). These two mutations were homozygous viable and showed no significant difference in embryo hatching rate when compared to control *w^1118^* wild-type flies (**Figure 1B**). The null nature of these mutations was confirmed by western blot (WB) and immunofluorescence (IF) analyses that showed no detectable dBigH1 in ovaries and early embryos from homozygous *bigH1^NULL^* and *bigH1^NSTOP^* flies (**Figures 1C** and **1D**). These results show that lack of *bigH1* does not significantly affect embryo viability, which is in agreement with Li *et al*. (2019).

**Figure 1.**
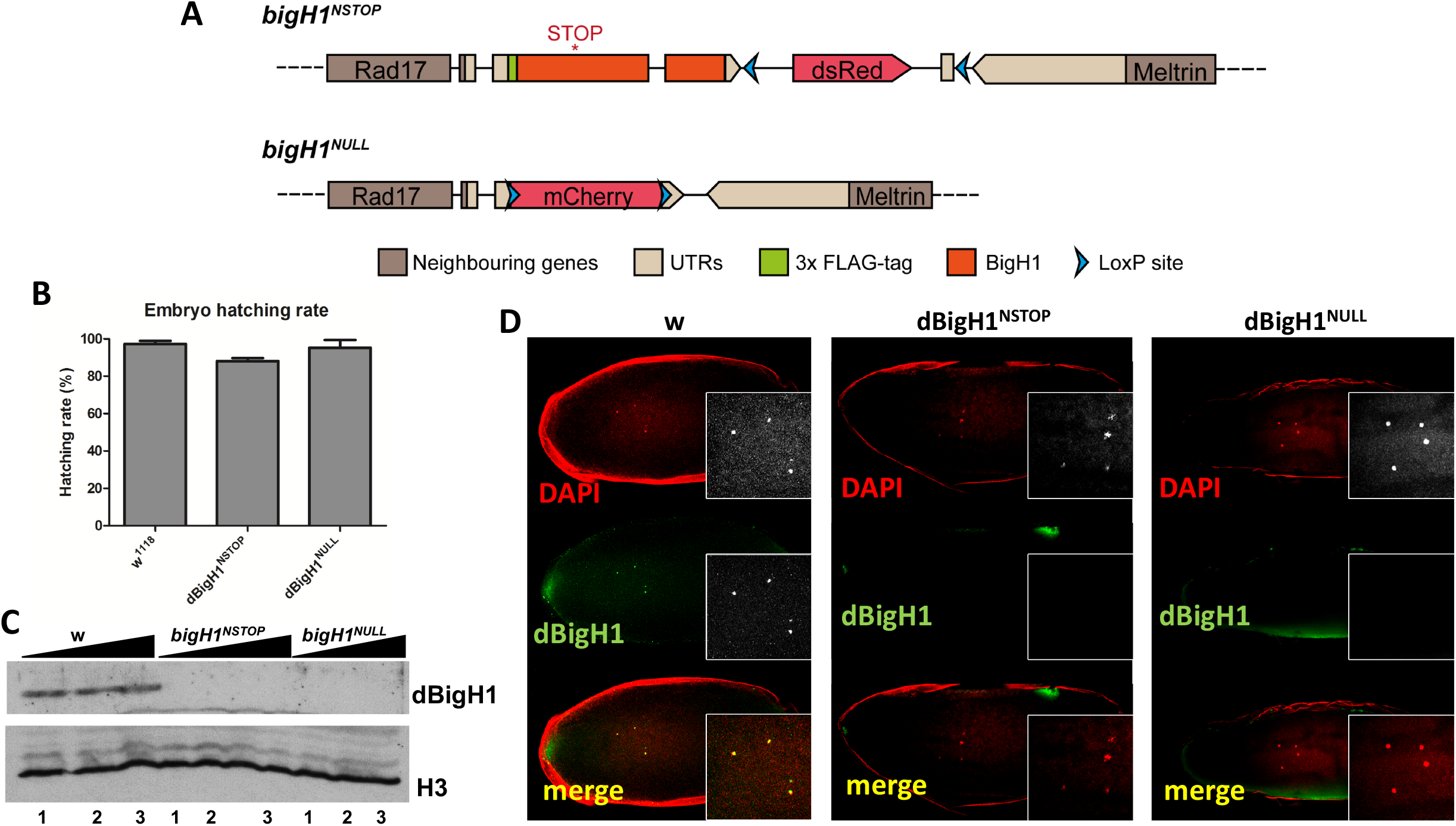
Generation of null *bigH1* CRISPR alleles. **A**) Schematic representation of the *bigH1^NULL^* and *bigH1^NSTOP^* alleles. **B**) The rate of embryo hatching is presented for the indicated genotypes. **C**) Western-blot analysis with αdBigH1 antibodies of increasing amounts (lanes 1-3) of total extract prepared from ovaries of females of the indicated genotypes. αH3 was used for loading control. **D**) Immunofluorescence staining with αdBigH1 antibodies (in green) of early *Drosophila* embryos (nc =2) of the indicated genotypes. DNA was stained with DAPI (in red).

Next, we analyzed whether, in null *bigH1^NULL^* and *bigH1^NSTOP^* early embryos, the lack of dBigH1 was compensated by the early expression of maternal dH1 as proposed in Li *et al*. (2019). Earlier to nuclear cycle (nc) 7, we could not detect αdH1 immunostaining in control wild-type embryos (**Figure 2A**). Instead, we observed intense αdH1 immunostaining in 100% of homozygous *bigH1^NULL^* (N= 40) and *bigH1^NSTOP^* (N= 30) embryos before nc 7 (**Figures 2C** and **2D**), confirming the compensatory dH1 expression described in Li *et al*. (2019).

**Figure 2.**
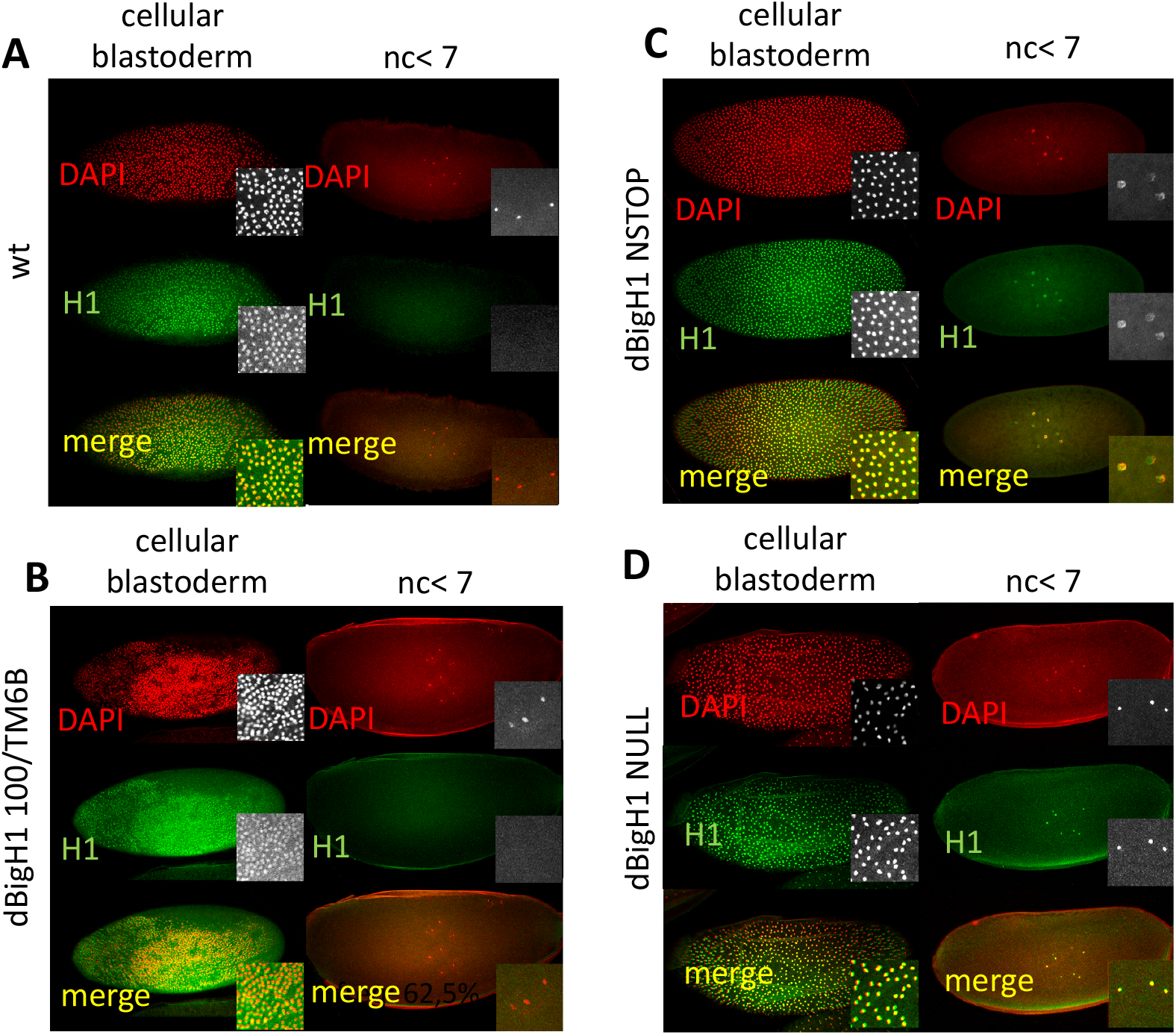
Compensatory dH1 expression. **A**) Immunofluorescence staining with αdH1 antibodies (in green) of wild-type *Drosophila* embryos at the cellular blastoderm stage (left) and before nc 7 (right). DNA was stained with DAPI (in red). Insets show enlarged images. **B**) As in **A** but for embryos from *bigH1^100^/TM6B* parents. **C**) As in **A** but for embryos from homozygous *bigH1^NSTOP^* parents. **D**) As in **A** but for embryos from homozygous *bigH1^NULL^* parents.

Results reported above and in Li *et al*. (2019) are in contrast with the high lethality observed in homozygous *bigH1^100^/bigH1^100^* embryos (Pérez-Montero et al., 2013). This lethality was observed in crosses between heterozygous *bigH1^100^/TM6B* parents, in which the maternal dBigH1 contribution was reduced but not abolished (Pérez-Montero et al., 2013). Thus, we wonder whether the compensatory dH1 expression was activated in these embryos. For this purpose, we performed IF experiments that failed to detect significant αdH1 signal in embryos before nc 7 (N=28) (**Figure 2B**), which is in contrast with the high reactivity detected in homozygous *bigH1^NULL^* and *bigH1^NSTOP^* mutant embryos (**Figures 2C** and **2D**). These results indicate that reducing the maternal dBigH1 contribution in *bigH1^100^/TM6B* flies is not sufficient to activate compensatory dH1 expression.

## DISCUSSION

Results presented here confirmed those previously reported in Li *et al*. (2019), showing that the lack of embryonic dBigH1 linker histone activates the compensatory expression of the somatic dH1 counterpart in early embryos, at stages in which dH1 is normally not expressed. As a consequence, embryos lacking dBigH1 show no detectable developmental defects and are viable. In contrast, embryos coming from heterozygous *bigH1^100^* mothers do not show compensatory dH1 expression, likely because their maternal dBigH1 content is sufficiently high. In this regard, the lethality of homozygous *bigH1^100^/bigH1^100^* embryos might indicate a failure to compensate their reduced maternal dBigH1 contribution with the expression of somatic dH1. However, we cannot exclude the possibility that a secondary mutation accumulated in the stock is causing this lethality. Other important questions remain equally unanswered. For instance, it would be interesting to determine whether the lack of any linker histone causes embryo lethality. Finally, the mechanism by which the lack of dBigH1 activates early expression of maternal dH1 also needs to be investigated.

## ACKNOWLEDGEMENTS

We are thankful to Dr. J. Kadonaga for the αdH1 antibodies. This work was supported by grants from the MICINN (PGC2018-094538-B-100) and the “Generalitat de Catalunya” (SGR2017-475) to FA, and the Hungarian National Scientific Research Fund (OTKA-116372) to IB and LH.

